# The effects of Polynucleotides-based biomimetic hydrogels in tissue repair: a 2D and 3D *in vitro* study

**DOI:** 10.1101/2025.03.18.643887

**Authors:** Maria Teresa Colangelo, Stefano Guizzardi, Luana Laschera, Marco Meleti, Carlo Galli

**Affiliations:** Department of Medicine and Surgery, Histology and Embryology Lab, University of Parma, Parma, Italy; Department of Medicine and Surgery, Dental School, University of Parma, Parma, Italy

**Keywords:** Polynucleotides, Fibroblasts, Wound healing, Cell movement, Cell shape

## Abstract

Biomimetics offer promising tools to improve wound healing in difficult clinical conditions. Polynucleotides (PN) show high potential for tissue repair in oral and periodontal surgery, by relying on the body’s inherent self-healing capabilities. The aim of the present study was to elucidate in vitro the effects of Odonto-PN(O-PN) and Regenfast (REG), two PN-based compounds, on oral tissue repair. We employed 3D spheroid cultures and cell scratch assays to simulate wound healing in vitro, assessing cell migration and morphology under normal conditions and following mitomycin-induced inhibition of cell growth. Both O-PN and REG promoted early cell viability and spheroid disassembly. O-PN supported initial outgrowth of fibroblasts, whereas REG enhanced sustained cell migration at later time points. In scratch assays, REG effectively facilitated defect closure—even under mitomycin treatment—and induced a more elongated, migratory cell phenotype. These findings suggest that both O-PN and REG can favorably modulate fibroblast function to support wound repair. While O-PN fosters early activation and cell viability, REG exerts potent promigratory effects that may be particularly useful for complex periodontal regeneration. Their selective use could provide valuable adjuncts in clinical protocols aimed at restoring delicate oral structures, such as the interdental papillae.

## INTRODUCTION

Wound healing is a physiological phenomenon crucial for repairing damaged tissues and restoring structural and functional integrity [1]. Although the human body has significant regenerative potential [2], there are numerous clinical scenarios that require targeted interventions to support this process [3,4]. Dentistry, in particular, faces unique regenerative challenges, including the repair of periodontal tissues and the regeneration of delicate structures such as gingiva and papillae, which demand specialized therapeutic approaches [5]. Addressing these challenges requires a deep understanding of the cellular and molecular mechanisms that drive wound healing [6]. This begins with the formation of a provisional matrix, which provides structural support and serves as a scaffold for incoming cells [7]. An inflammatory phase then initiates, clearing out debris and pathogens, and setting the stage for subsequent tissue repair [8]. Cellular precursors from neighboring tissues proliferate and migrate into the matrix [9], to deposit new sound tissue [10]. In cases where the body’s natural processes are insufficient, innovative therapeutic approaches become essential. Ideally, tailored therapies should address specific impairments in the healing process to counteract any defective mechanisms that may lead to incomplete healing, based on the patient’s individual clinical needs [11].

To meet the challenges of incomplete natural healing, biomimetic strategies have emerged as promising tools to enhance wound care [12–15] for their ability to emulate natural tissue components and replicate biological mechanisms supporting physiological healing. By imitating the structure and function of biological materials, biomimicry offers therapeutic solutions that align closely with the body’s own repair mechanisms. Importantly, the mechanical properties of these biomimetic substrates—e.g. their viscoelasticity—play a pivotal role in controlling cell responses. The substrate’s physical characteristics, including its stiffness and elasticity, influence cell adhesion, migration, and activity, thereby creating a microenvironment that actively supports the healing process [16]. Extensive research has demonstrated that by adjusting these physical properties in scaffolds to mimic those of native tissues, it is possible to guide cellular responses in desired directions, enhancing tissue repair and regeneration [17–22].

A noteworthy implementation of this paradigm can be found in the application of polynucleotides or hyaluronic acid for wound healing [23,24]. Polynucleotides (PN) are a DNA derivative of fish origin [25]. PN fill intercellular spaces and act as a supportive extracellular matrix. This matrix not only provides essential hydration for volume maintenance and thus spatial integrity but also establishes a scaffold that fosters cell adhesion and communication. By integrating within the extracellular environment, PN contribute to a stable microenvironment that mimics physiological conditions, which is crucial for the natural tissue regeneration process. This hydrated and structured support encourages cellular migration and alignment, enhancing the extracellular matrix and promoting tissue resilience [24]. Extensive research has underscored the versatility of PN, showcasing their effectiveness in vitro [26]. Moreover, there is an abundant literature about their successful application in diverse in vivo contexts [27–30].

In the present research, we investigated two PN-based compounds for use in dentistry, Odonto-PN and Regenfast; the former contains exclusively PN, while the latter contains PN associated to hyaluronic acid (HA), a highly biocompatible molecule abundantly distributed in human tissues and with a vast literature supporting its use as a hydrogel scaffold in numerous approaches [31–33]. Building on these properties, the aim of this study is to investigate how these compounds support tissue regeneration by discriminating cell growth and cell migration, using an in vitro model of primary human gingival fibroblasts.

## MATERIALS AND METHODS

### Materials

The products used for the present study are Class III medical devices Odonto-PN (20 mg/mL) and Regenfast (10 mg/mL PN and 10 mg/mL HA), both manufactured by Mastelli s.r.l., Sanremo, Italy. Polynucleotides (PN) from Mastelli s.r.l., Sanremo, Italy, is a compound holding DNA of different chain lengths obtained from salmon trout (Oncorhynchus mykiss) gonads through a proprietary high purification technology (HPT^TM^), which provide high-quality DNA without immunological side effects [34].

### Cell Culture

The cell model used was HGF (Primary Gingival Fibroblast; Normal, Human, Adult) from ATCC (LGC Standards S.R.L., Milan, Italy) and cultured in complete Dulbecco modified MEM (DMEM, LifeTechnologies, Carlsbad, CA, USA) with 10% Fetal Bovine Serum (FBS, LifeTechnologies), 4 mM L-glutamine (Merck KGaA, Darmstadt, Germany), 100 IU/mL penicillin and 100 μg/mL streptomycin (PenStrep, Merck KGaA), in a humidified atmosphere at 37 °C and 5% CO_2_. Nikon TMS inverted optical microscope (Nikon, Tokyo, Japan) in phase contrast was used to follow up cell culture and a Nikon Digital Sight DS-2Mv acquisition system and NIS Elements F software (Nikon) were used for image analysis. For all experiments, fibroblasts were used from 4^th^ to 10^th^ passage. For each experiment, 3 groups were evaluated: Control, Odonto-PN (O-PN), and Regenfast (REG). Cells were exposed to 100 µg/ml O-PN or REG 24h after seeding; culture medium was used as control. The concentration of the compounds was based on previous literature [37].

### Morphometric analysis

Spheroids were obtained by seeding 6 × 10^3^ cells/well in 96-well plates specifically designed for 3D cultures (BIOFLOAT™ 96-well U-bottom Plates, faCellitate, Mannheim Germany), with 12 replicates for each condition. Twenty-four hours after seeding, cells were exposed to the compounds. To assess the effect of the compounds on cell viability and/or migration, the spheroids were re-plated on a standard 96-well culture plate after a week of culture and maintained under routine conditions for 3 days. The 3D culture images were collected over time by an inverted optical microscope equipped with a camera. The shape of the spheroids, and the presence and position of adhering cells on the plastic culture around the spheroids were evaluated (ImageJ, U.S National Institutes of Health, Bethesda, MD, USA) by a blinded observer.

### Cell growth

Cells were seeded in a 24-well plate at a density of 3x10^4^ cells/well, with 6 replicates for each condition. Twenty-four hours after seeding, fibroblasts were treated with Mitomycin C (10µg/ml) for 4 hours and then exposed to O-PN, REG or culture medium. Samples exposed to O-PN, REG or culture medium in the absence of Mitomycin C were used as controls. Cell number was evaluated after 72 h of exposure with Scepter 3.0 Handland Automated Cell Counter (Sigma Aldrich; Merck KGaA).

### In Vitro Scratch Assay

Briefly, 15 × 10^4^ cells were seeded into a 12-well plate, in triplicate, and, upon confluence, were treated with Mitomycin C (10 µg/ml) for 4 hours to inhibit cell growth. The cell monolayer was then scratched with a 200 μL plastic pipette tip across the center of the well and cells were exposed to O-PN, REG or culture medium. Samples not pretreated with Mitomycin C and exposed to O-PN, REG, or culture medium were used as controls. We monitored the closure of the scratch for up to 72 h after treatment and acquired microphotographs over time. At 72h, samples were fixed and stained with Giemsa solution. The acquired images were used to perform a morphometric analysis with ImageJ.

### Statistical Analysis

Data analysis was performed using Prism X (GraphPad, La Jolla, CA, USA). Data were expressed as means ± standard deviation (mean ± SD) of three repeated experiments. We investigated differences between the experimental group using two-way ANOVA test with Bonferroni post-test, and p-value < 0.05 was considered a statistically significant difference.

## RESULTS

### REG promote cell migration from the spheroids

We cultured cells as spheroids on special anti-adhesion plates and added vehicle, O-PN, or REG. The spheroids were then transferred to regular culture multi-well plates (Fig. 1).

**Figure 1.**
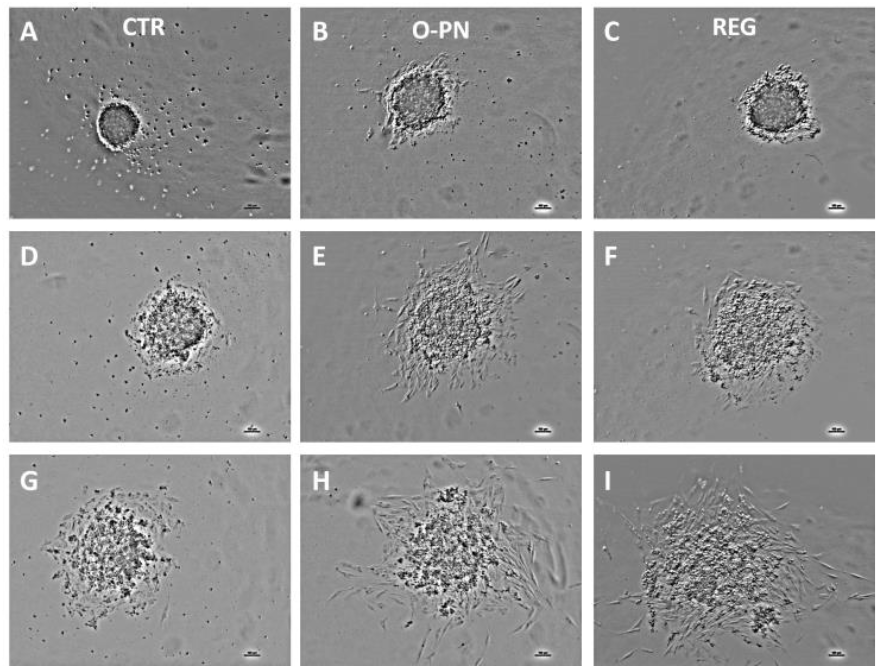
Phase-contrast microphotographs of HGF spheroids previously exposed to culture medium (A, D, G), O-PN (B, E, H) or REG (C, F, I) for 1 week and observed 24 (A-C), 48 (D-F) or 72 hours (G-I) after being replated in regular cell culture plates. (Bar= 100 µm, Magnification=10X)

We monitored the plates to observe cells moving out of spheroids and across the plate in a growth/migration process. In vehicle-treated samples, spheroids maintained their integrity 24 hours after replating (Fig. 1A), with only a few round cells lying scattered around the spheroids. These cells were likely dead and had detached during the transfer, settling in the surrounding area. Forty-eight hours after re-plating, the spheroids appeared unchanged or slightly larger (Fig. 1D), occasionally appearing flatter, with more irregular margins as cells began to migrate outward. By 72 hours, visible signs of cell migration were evident, with cells forming a thin halo around the spheroids (Fig. 1G), further increasing the irregularity of the spheroid perimeter at this time point.

Adding O-PN to the spheroids prior to re-plating resulted in viable cells visible around the spheroids already 24 hours after re-plating (Fig. 1B). While it’s possible that these cells migrated out of the spheroids soon after re-plating, their presence may also be due to cells detaching during the transfer; notably, they appeared viable based on firm attachment to the substrate and the absence of morphological signs of distress. By 48 and 72 hours after re-plating, growth and/or migration in O-PN samples became extensive (Fig. 1E, 1H). The spheroids appeared flattened and more irregular in shape, with migrating cells forming columns that extended outward in multiple directions (Fig. 1H).

Cell behavior appeared similar in REG samples at 24 and 48 hours of culture after re-plating (Fig. 1C, 1F); here adhering cells could be observed lying closely around the spheroids. However, by 72 hours, cells in REG samples had proliferated/migrated across a larger area than in O-PN-treated samples (Fig. 1I), the spheroids started to significantly dissolve, indicating that more cells from the spheroids were dispersing onto the plate.

We then measured how far cells migrated under different conditions, as the distance between the approximate center of the spheroid and the farthest cell on the plate (Fig. 2A). As expected, this distance increased over time in every group (Fig. 2B). After 72 hours, cells in REG samples had migrated more than 900 micrometers from the spheroids, compared to an average distance of less than 600 micrometers in control samples—a statistically significant difference (Fig. 2B).

**Figure 2.**
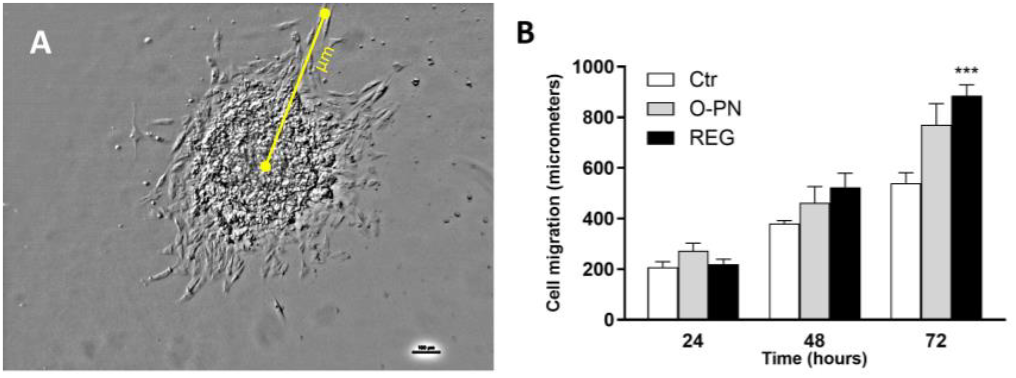
A) Phase-contrast microphotograph of HGF spheroid replated in a regular culture plate. Maximum cell migration was measured as the distance between the spheroid center and the farthest cell (yellow line). B) Bar chart showing the maximum cell migration of HGFs from spheroids 24, 48, or 72 hours after replating. ***=p<0.001 vs control.

### REG increases cell migration in wound healing cell assay

While the spheroid re-plating model was quite effective at providing visible and measurable signs of cell migration, to isolate the effects of the tested compounds on cell movement, we inhibited cell growth using mitomycin. Figure 3 shows the effect of O-PN or REG on fibroblasts in the absence or in the presence of mitomycin after 72 hours of culture. When growth was unimpeded (left side of the graph), the experimental compounds significantly increased the number of cells per well compared to the untreated control, with REG samples showing a significantly higher cell count. However, mitomycin eliminated any differences across the groups, indicating that it had completely suppressed the effects of the compounds on cell numbers.

**Figure 3.**
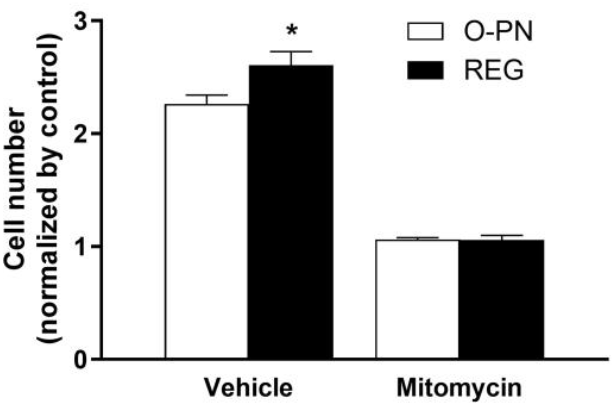
Bar chart of HGF number in the absence or in the presence of mitomycin and after treatment with O-PN or REG for 72 hours. All values were normalized by the control group stimulated with culture medium. *= p<0.05 vs O-PN.

Cell scratch assays are *in vitro* assays where a scratch is created on the surface of a confluent cell monolayer. By observing how cells recolonize the gap from the sides, the assay provides a simplified model for evaluating wound healing. The healing process naturally involves two main events that are recapitulated in this model, cell growth, once contact inhibition is removed after the scratch is created, and cell migration into the defect. To better distinguish the relative contributions of growth and migration, we conducted this assay with and without mitomycin, in the presence of O-PN or REG (Fig. 4). Microphotographs in the left column (Fig. 4A, C, E) show cells repopulating the defect area after 72 hours, either untreated (control) or exposed to the compounds without mitomycin. Cells within the defect were colorized with ImageJ to improve visibility. Visual inspection suggested a higher cell density (and a narrower defect) in the REG group compared to control samples, confirmed by quantification of occupied area through image analysis (Fig. 4G). In control samples, only 30% of the defect area was occupied by cells—significantly less than in the experimental samples (Fig. 4G, lefthand side of the graph). Additionally, the defect appeared narrower in REG sample (Fig. 4E) than in control samples (Fig. 4A), a finding confirmed also by image analysis (Fig. 4H).

**Figure 4.**
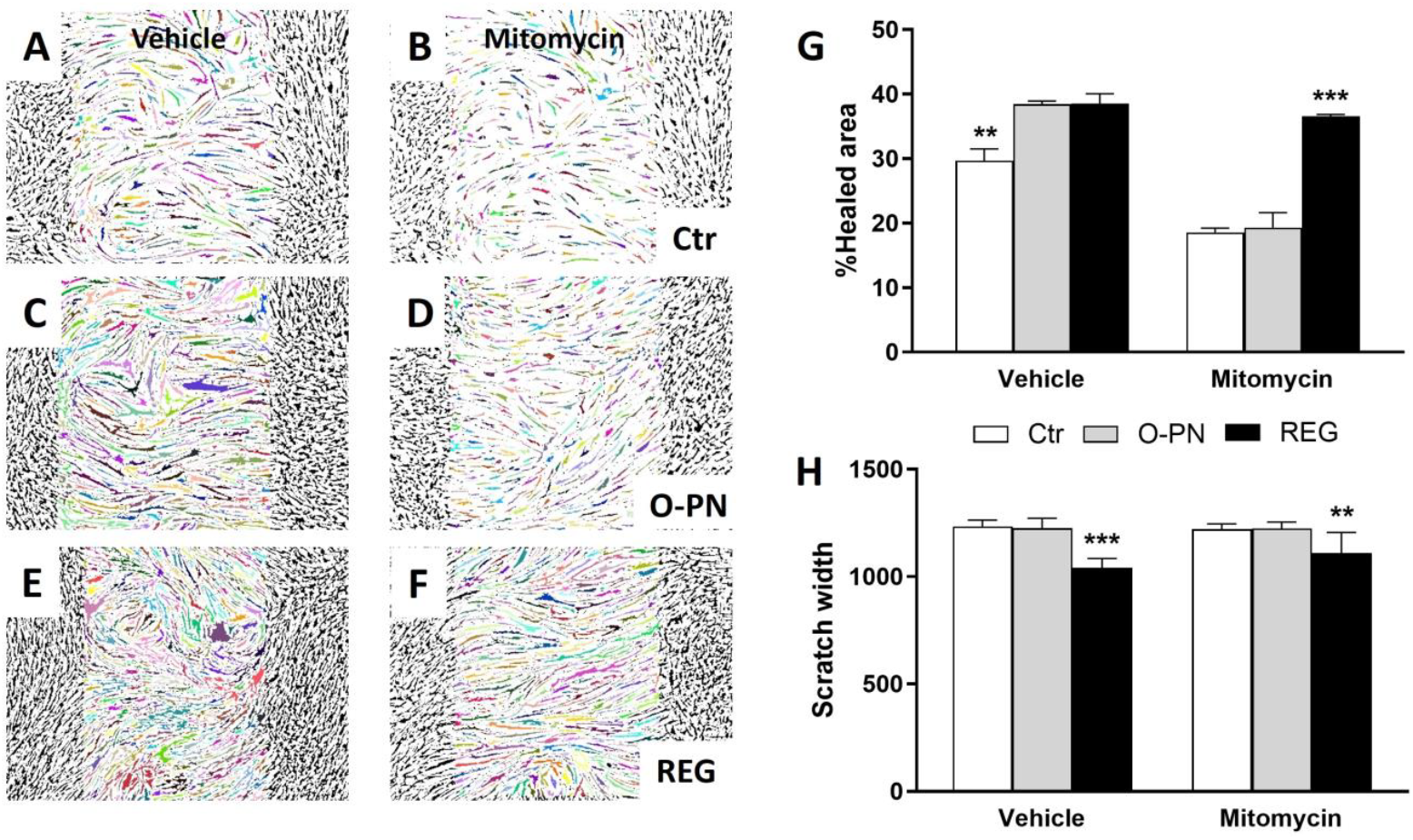
A-F) ImageJ masks of scratch assay after 72 hours of healing in the absence (A, C, E) or in the presence (B, D, F) of mitomycin and after treatment with medium (A, B), O-PN (C, D), or REG (E, F). Images were colorized to highlight cells repopulating the defects. G) Scratch area covered with cells and scratch width in the presence or in the absence of mitomycin and after stimulation with medium, O-PN, or REG for 72 hours was quantitated with ImageJ and is illustrated as bar charts; **= p<0.001 vs O-PN, or REG; ***=p<0.001 vs Ctr, O-PN. H) Scratch width in the presence or in the absence of mitomycin and after treatment with experimental compounds for 72 hours and expressed as bar chart; ***=p<0.001 vs Ctr, O-PN; **= p<0.01 vs Ctr, O-PN.

Adding mitomycin visibly reduced the occupied area at 72 hours, bringing it down to about 20% in control samples and in those exposed to O-PN alone (Fig. 4G, right hand side of the graph). This suggests that the effect of PN alone on wound healing primarily depended on promoting cell growth. However, in the presence of mitomycin, the occupied area in REG samples remained significantly higher than in other groups (Fig. 4G), indicating that REG effectively supported cell migration to fill the defect. Similarly, defects exposed to REG were still narrower than in the control or O-PN group even after addition of mitomycin (Fig. 4H).

### REG treatment induces a more elongated phenotype in cells

Figure 5 shows high-magnification (20x) images of representative fields from the defect area in control samples and samples exposed to O-PN or REG, all in the presence of mitomycin. Visual inspection revealed that cells in the control group (Fig. 5A, D, G) displayed heterogeneous morphology. Some cells appeared larger, with a wide cytoplasm and thick pseudopods anchoring them to the substrate, while other cells were narrower, with the nuclei protruding from an elongated, thin cytoplasm. Cell orientation was also varied, with cells aligned in different directions within the same field. Cells in treated samples appeared more elongated than in the control group, but with noticeable differences. Cells in the O-PN group (Fig. 5B, E, H), and even more so in the REG group (Fig. 5C, F, I) mostly conformed to a very thin, spindle like and elongated morphology with neighboring cells aligned in the same direction.

**Figure 5.**
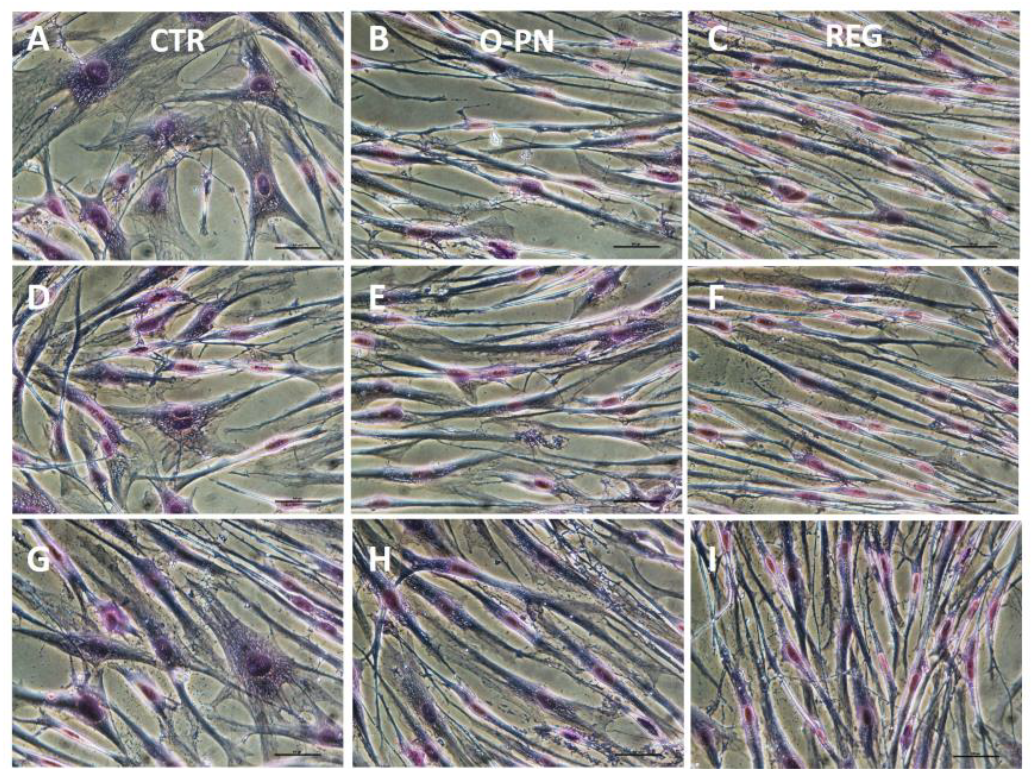
Phase-contrast microphotographs of HGF treated with mitomycin and with culture medium (A, D, G), O-PN (B, E, H) or REG (C, F, I) for 72 hours. (Bar= 50 µm, Magnification=20X). Cells were stained with Giemsa.

This was confirmed through cell circularity analysis by ImageJ (Fig. 6). Cell circularity is an index calculated as the ratio between a cell’s main axes, ranging from 0 to 1: a value of 1 indicates a perfect circle (equal diameters), while values closer to 0 indicate a more elongated, line-like shape. Without mitomycin cells in the O-PN or REG groups were significantly more elongated than in the control group. The addition of mitomycin did not affect cell shape in the REG group, though cells in the other groups tended to become more circular. Consequently, in the presence of mitomycin, cells in the REG group had significantly lower circularity values than those in the other groups, and cells in the O-PN group had lower circularity than the control group.

**Figure 6.**
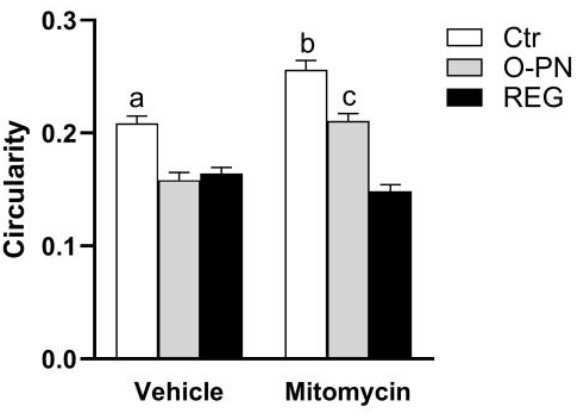
Bar graph of the circularity coefficient for HGF cells in the absence or in the presence of mitomycin and added with medium, O-PN or REG, for 72 hours. a, b=p<0.001 vs O-PN, REG; c =p<0.001 vs Ctr, REG.

## DISCUSSION

Wound healing is a complex process that requires a coordinated series of events to restore damaged tissue [6]. These include cell activation and proliferation and the formation of a provisional matrix [16], which matures over time, allowing the tissue to regain its structural integrity and function. However, complex clinical scenarios, such as periodontal repairs, often require additional therapeutic strategies to ensure successful outcomes [35]. In particular, regenerating interdental structures like the papilla is challenging due to limited blood supply and tissue volume, which restricts cellular migration and matrix formation [36]. In such situations, adjunctive therapies that support tissue trophism can make a critical difference in restoring aesthetics and functionality.

Given these challenges, polynucleotides (PN) have shown potential in clinical practice for tissue repair, demonstrating benefits in cell growth and viability across various studies. Evidence from in vitro and in vivo studies indicates that compounds based on polynucleotides increase cell growth and support cell viability [24–26,37]. Cell migration represents a necessary and dynamic event in wound healing, driven by internal and external forces [38], generated by the cytoskeleton and by cell-substrate interactions [39]. Based on the idea that hyaluronic acid is a common component of the extracellular matrix [40], i.e. the extracellular environment that provides anchoring and biological cues that direct cell activity [41], including cell migration [42], we set out to investigate how O-PN and REG could differentially control cell growth and motility.

In a 3D spheroid model, our results showed that O-PN initiated an early activation of cells, well noticeable by 48 hours, as evidenced by a thicker halo of cells around the spheroids. This suggests that O-PN may help cells transition out of a quiescent or compact state, supporting initial fibroblast spreading and viability. By 72 hours, however, spheroids in the REG group showed more extensive disassembly and a larger halo of migrating cells, indicating that REG provides robust cues for cell motility.

Measuring the distance between the spheroid center and the farthest migrating cell also highlighted the stronger migratory response in the REG group. While the apparent edge of REG in motivating cell migration was clear, O-PN also promoted noticeable cell activation. Thus, O-PN may be beneficial in kick-starting early cell outgrowth and in creating a scaffolding microenvironment that maintains cell viability, while REG can be effective for sustained migration. However, this assay had some unavoidable limitations. Although two operators (C.G. and M.T.C.) reached a consensus on the spheroid center and the farthest cell for each field, a degree of subjectivity in these measurements could introduce a margin of error. Additionally, while the centroid was used as the starting point for distance measurement, it is likely that cells at the periphery migrated from the spheroid edge rather than its center, leading to a possible overestimation of migration (though comparisons across treatment groups remain valid). Finally, part of the cell population around the spheroid may have reached its positions through growth, not migration alone; thus, cell placement likely reflects a combination of growth and migration. Although the 72-hour observation period likely limited growth to a maximum of two cycles, any treatment effects on cell growth could influence the final positions of cells around the spheroid.

To further distinguish the role of migration from growth, we used mitomycin to inhibit cell growth [43], eliminating any contribution from the tested compounds (Fig. 3). We then proceeded to investigate to what extent presence of O-PN or REG would increase scratch healing. We assumed that mitomycin would impair the closure of cell scratches because cells would not proliferate and thus there would be fewer cells available to fill the gap. This effect was observed in the control and O-PN samples (Fig. 4B, D, G), but not in the REG group (Fig. 4F, G), where the occupied (‘healed’) defect area remained unchanged even after mitomycin. However, this does not imply that the healing process in the REG group was unaffected; visual inspection of the defect revealed differences in cell morphology. Our image analysis does not distinguish individual cells, so colored areas could consist of individual cells or cell clusters. The elongated shape of these elements likely explains their ability to cover wider areas of the substrate, even though mitomycin prevented growth, thus limiting the number of cells available to migrate from the defect margin. Figure 5 confirms that individual cells were longer and thinner in the REG, and, to a lesser extent, in the O-PN groups after addition of mitomycin. This elongated shape is associated with cell migration [44], as cells extend pseudopods to move across the culture plate [45]. Under the experimental conditions of mitomycin-induced growth inhibition, however, REG created a scaffold that was supportive enough for effective cell migration to attain a better defect coverage, which might prove useful in situations where wound closure must be effectively achieved, such as in graft surgery [46]. Given the strong connection between cell morphology, the cytoskeleton, and cellular adhesion, the cell scratch assay reveals a potential synergistic effect of PN and HA in REG. Although the precise mechanisms underlying this enhanced cell migration remain still unclear, the combination of these components may have created a scaffold network that provided effective adhesion points, through the optimal level of viscoelasticity to stimulate cell activity and facilitate movement of gingival fibroblasts [7,19,47]. On the other hand, O-PN appears to excel in supporting cell viability and could be an optimal choice in conditions where maintaining adequate tissue trophism is necessary even under physiological or anatomical constraints, such as in papilla regeneration.

Future investigations should explore these differences to inform tailored clinical protocols, where the choice of O-PN or REG could be based on the specific healing requirements of a given defect or patient population.

Overall, our data confirm that polynucleotide-based compounds through mechanical scaffold action can significantly support viability and motility of human gingival fibroblasts, key players in periodontal wound repair.

Notably, O-PN appeared to activate cells earlier in the assay, while REG significantly enhanced cell motility in culture. In an in vitro wound healing model, we found that both O-PN and REG support wound closure. In the case of REG, cell migration into the defect was promoted to the point that it could support effective defect healing even after blocking cell growth.

These findings highlight the potential of O-PN and REG to promote cell growth and activation, and in the case of REG of expediting cell migration in situations of wound closure.

## Acknowledgements

The authors would like to thank dr. Silvana Belletti for her advice.

## Author contribution statement

Conceptualization, C.G. and S.G.; methodology, L.L. and M.T.C.; formal analysis, C.G.; resources, S.G. and M.M.; writing—original draft preparation, C.G.; writing— review and editing, M.M.; visualization, C.G.; supervision, S.G. All authors have read and agreed to the published version of the manuscript.

